# Climatic sensitivity of species’ vegetative and reproductive phenology in a Hawaiian montane wet forest

**DOI:** 10.1101/749614

**Authors:** Stephanie Pau, Susan Cordell, Rebecca Ostertag, Lawren Sack, Faith Inman-Narahari

## Abstract

Understanding the way tropical tree phenology (i.e., the timing and amount of seed and leaf production) responds to climate is vital for predicting how climate change may alter ecological functioning of tropical forests. We examined the effects of temperature, rainfall, and photosynthetically active radiation (PAR) on seed and leaf phenology in a montane wet forest on Hawaiʻi using monthly data collected over ∼6 years. We expected that species’ phenologies were more sensitive to temperature and PAR than to rainfall at this wet tropical site because rainfall is not limiting. Seed production declined with increasing temperatures for two foundational species in Hawaiian forests (*Acacia koa* and *Metrosideros polymorpha*). Seed production also declined with rainfall for two species, and greater PAR for one species. One species showed relatively flat responses to climate. Community-level leaf phenology was not strongly seasonal. Unlike seed phenology, we found no effect of temperature on leaf phenology. However, leaf fall increased with rainfall. Climatic factors explained a low to moderate proportion of variance for both seed and leaf litterfall, thus the impact of future climate change on this forest will depend on how climate change interacts with other factors such as daylength, biotic, and/or evolutionary constraints. Our results nonetheless provide insight into how climate change may differentially affect different species with potential consequences for shifts in species distributions and community composition.

---

There is little information on changes in tropical plant phenology in response to climate change, unlike the mounting evidence from temperate regions for “spring advancement”, i.e., the earlier occurrence of first bud burst and flowering due to warming temperatures (Menzel *et al*. 2006, Cook *et al*. 2012, Abernethy *et al*. 2018). Seasonal changes in temperate regions are more pronounced than in the tropics, thus the phenological sensitivities of temperate species to climate change are easier to identify (Newstrom *et al*. 1994, Pau *et al*. 2011, Cook *et al*. 2012). In the tropics, the growing season is potentially year-round. Consequently tropical plant phenology has been hypothesized to be more closely tied to biotic factors such as pollinator abundance or competition for resources rather than seasonal changes in climate (Pau *et al*. 2011). Yet underlying selective forces vs. proximate factors that cue phenological events often interact and can be difficult to separate (Rathcke & Lacey 1985, van Schaik *et al*. 1993). Phenological cues are usually consistent changes in the abiotic environment, which then trigger physiological mechanisms controlling flower, fruit, or leaf production (van Schaik *et al*. 1993).

Because there is little variation in temperature in low latitudes, research on tropical plant phenology – both leaf and reproductive phenology – has focused on changes in rainfall (Frankie *et al*. 1974, Lieberman 1982, Wright 1991, Borchert 1994, Reich 1995, Zimmerman *et al*. 2007, Sakai & Kitajima 2019). Many tropical and subtropical regions experience a dry season associated with the seasonal movement of the Intertropical Convergence Zone (ITCZ). Although the dry season results in seasonal water deficits, there are also fewer clouds allowing greater light interception by the canopy. Both experiments and observational data show that greater light availability is associated with greater community-level flower, fruit, and leaf production in tropical forests (Graham *et al*. 2003, Wright & Calderón 2006, Zimmerman *et al*. 2007, Pau *et al*. 2013).

Variability in temperature is not as great in the tropics compared to temperate or high latitude regions, thus tropical species may be more sensitive to warming because these organisms are adapted to a narrower range of temperatures and may be living closer to their upper thermal limits (Janzen 1988, Tewksbury *et al*. 2008, Wright *et al*. 2009). Tropical regions are not warming as much or as fast as high latitude regions (IPCC 2014), yet the tropics may experience novel climates outside of their historical range much sooner (Williams *et al*. 2007, Mahlstein *et al*. 2011, Mora *et al*. 2013, 2015). Consequently the physiological tolerance of tropical species combined with the pace of environmental change will determine their vulnerability to climate change (Tewksbury *et al*. 2008, Kingsolver 2009).

Although temperature has not received much attention in the search for abiotic drivers of tropical phenology, temperature is a fundamental constraint on numerous biological processes (Kingsolver 2009). Seasonal flowering patterns were linked to changes in temperature at two tropical sites with long-term data (Pau *et al*. 2013). In a lowland moist seasonal forest in Panama and a montane ever-wet forest in Puerto Rico, warmer months were associated with greater flowering activity (Wright & Calderón 2006). There are few physiological experiments that have examined the temperature sensitivity of tropical reproduction, but reproductive organs are known to be highly sensitive to temperature (Larcher & Winter 1981, Slot & Winter 2016). It is unclear if tropical leaf phenology is also sensitive to temperature fluctuations.

There is remarkable diversity in patterns of tropical flowering (Newstrom *et al*. 1994, Sakai 2001), seed and leaf production (Frankie *et al*. 1974; Reich 1995; Leishman 2000). Identifying species’ phenological strategies should help us predict the potential ‘winners’ and ‘losers’ of climate change. For example, species that continuously reproduce throughout the year may be less strongly cued by climate and may thus be less sensitive to future climate change (Pau *et al*. 2011). The divergent responses of species’ reproductive phenologies to climate change may alter plant community composition if species are recruitment limited (Drake *et al*. 1998, Hubbell *et al*. 1999, Inman-Narahari *et al*. 2013). Plant phenology also has important cascading effects throughout the community by structuring the timing of food availability for many organisms.

The Hawaiian flora provides unique insight into the ecology and evolution of plant diversity, and is considered a model system due to its isolation, endemism and the relative simplicity of its processes due to low species diversity (Sakai *et al*. 1995, Wagner & Funk 1995, Price & Wagner 2004). Yet there are few published studies on the reproductive phenology of Hawaiian forests (e.g., van Riper III, 1980; Drake, 1992; Berlin *et al*., 2000). More work has examined monthly leaf litterfall, yet these studies have generally focused on leaf litter’s role in nutrient cycling and contributions to aboveground net primary productivity (e.g., Vitousek *et al*. 1995, Raich 1998, Schuur & Matson 2001, Austin 2002), not seasonality and responses to climatic variability. This lack of understanding limits our knowledge of how climate change may alter Hawaiian forest phenology and associated ecosystem functions. Hawaiian forests have lower tree diversity, but are structurally similar to other continental tropical forests (Ostertag *et al*. 2014) with tropical phenological strategies represented. Their extreme isolation in the Pacific makes them a unique signal for climate change impacts on ecological communities, unlike forests such as the Amazon, which experience large local feedbacks between the canopy and atmosphere (Kooperman *et al*. 2018), complicating the climate signal. In this study we examine monthly seed production of the four dominant tree species and community-wide leaf litterfall from a montane wet forest on the Island of Hawaiʻi, and compare their sensitivities (i.e., direction and magnitude of response) to temperature, rainfall, and photosynthetically active radiation (PAR). We hypothesize that species’ phenologies are more sensitive to temperature and PAR than to rainfall because tropical species should be especially sensitive to temperature and rainfall at this site is not limiting.

## METHODS

### study site, phenology, and climate data

The Laupāhoehoe Forest Dynamic Plot (FDP; 19°55’ N, 155°17’ W), part of the Forest Global Earth Observatory (ForestGEO) network (https://forestgeo.si.edu/), is a montane wet forest located at 1120 m in elevation on the Island of Hawaiʻi. The forest is comprised of 18 woody flowering tree species, 3 tree fern species, and the vegetation is highly representative of montane wet forests in Hawaiʻi (see Ostertag *et al*., 2014 for detailed site information). Mean annual precipitation is 3440 mm and mean annual temperature is 16° C (Giambelluca *et al*. 2013).

To monitor reproductive and leaf phenology, sixty-four litter traps within a 4-hectare plot were placed in a regular grid, 10-meters apart, and monitored as part of the Hawaiʻi Permanent Plot Network (HIPPNET). Reproductive (fruit and seeds) and leaf litterfall were censused each month following standard protocol (Wright *et al*. 2005). Fruits and seeds were identified to species and converted to number of seeds for each fruit, then summed across all traps for each month. Leaf litterfall, for all species combined, was summed across all traps, divided by the number of traps censused (because some months, traps were knocked over), and weighed each month. Sixty-five months of fruit/seed data were available between the months of October 2009 to March 2018 (with occasional missing collections between 2009 and 2015, only part of 2017 collected, and none of 2016 collected; a full 12-month record was only available for two years prohibiting year-to-year models). Thirty-two months of community leaf litterfall data were available. Because some collections could only be attributed to a month and year, leaf litterfall rates (e.g., g m^−2^ day^−1^) could not be determined accurately.

Climate stations at both sites are maintained by HIPPNET and record daily temperature (° C; HMP45C-L,Vaisala), rainfall (mm; tipping bucket rain gauge; TB3 CS700, Hydrological Services), and photosynthetically active radiation (PAR; μmol s^−1^m^−2^; Quantum sensor). All climate data were aggregated to monthly averages except for rainfall, which was summed each month.

Of the 21 plant species present at Laupāhoehoe, 12 were present at least once in the litter baskets and 4 dominant species were ultimately examined: *Metrosideros polymorpha* (’ōhi’a lehua; bird or insect pollinated, wind dispersed, Myrtaceae), *Acacia koa* (koa; insect pollinated, wind dispersed, Fabaceae), *Coprosma rhynchocarpa* (pilo; wind pollinated, bird dispersed, Rubiaceae), and *Cheirodendron trigynum* (ʻōlapa; bird or insect pollinated, bird dispersed, Araliaceae). These four dominant species comprise 57.5% of the relative abundance and 52.7% of the total basal area of the Laupāhoehoe FDP and were the trees that reached canopy and sub-canopy levels (Ostertag *et al*. 2014). The other 8 species occurred too infrequently in the litter traps for statistical analysis.

### statisical analyses

To examine relationships between seed production (counts) and leaf litterfall (grams) with climatic predictors, we used generalized additive models (GAMs) to estimate flexible (potentially non-linear) and independent smoothing functions to predictors and response variables (Venables & Ripley 1999, Zuur *et al*. 2009). We used a Poisson log-link likelihood for seed production and a Gaussian log-link likelihood for leaf litterfall, and estimated relationships between seed production or leaf litterfall and climatic predictors in GAMs using a cubic regression spline. Using GAMs we estimated response curves (i.e., ‘smooth terms’) to climatic predictors, temperature, rainfall, and PAR as well as month (to account for monthly seasonality separate from climatic seasonality) and year (to account for any yearly trend). We also included daylength as a predictor, but large (> 0.80) concurvity values (i.e., a generalization of collinearity for smooth terms in GAMs) limited accurate interpretation (see Supporting Information; Fig. S1 and S2, Table S1). Because time-series data are often non-independent, we accounted for serial autocorrelation in the error term using an AR(1), i.e., an autoregressive term of 1 month. Model residuals were examined and showed no significant autocorrelation. Seed production was modeled separately for each species using concurrent monthly climate data because fruit maturation, dispersal, and germination should be timed to climate (van Schaik *et al*. 1993). Leaf litterfall, however, is not as clearly timed to climate as leaf production might be because leaves are longer lived on the canopy. Thus leaf litterfall was examined using collection dates lagged one month prior to collection (when leaves had not yet fallen and are still on the canopy) because Pearson’s correlation coefficients were stronger with one-month lag compared to a two- or three-month lag.

In addition to examining each species’ seed production and community-wide leaf litterfall responses to light, temperature, and rainfall (i.e., the full model), we compared all possible reduced models using the Akaike Information Criterion (AIC) (Burnham & Anderson 2010) to assess which combination of climatic factors resulted in the best-fit model.

## RESULTS

### seed production

The most strongly seasonal species was *C. rhynchocarpa*, with protracted annual seed production from June-February and peak seed production in December (Figure 1). The other three species generally produced seeds year-round however some still exhibited seasonality in seed production. *C. trigynum* had a clear peak season of seed production from July-November, whereas seed production by *A. koa* and *M. polymorpha* were highly variable throughout the year*. A. koa* had the most variable monthly seed production with a coefficient of variation (CV) ranging each year from 0.85 – 3.00 (examining only 5 years where every month was represented). The CV of *C. trigynum* ranged from 0.65 – 1.35, *M. polymorpha* ranged from 0.77 – 1.18, and *C. rhynchocarpa* ranged from 0.22 – 1.05. Across years, *A. koa* seed production had the largest interannual variation indicated by the highest CV (1.24), followed by *C. rhynchocarpa* (0.74), *C. trigynum* (0.64), *M. polymorpha* (0.38).

**Figure 1.**
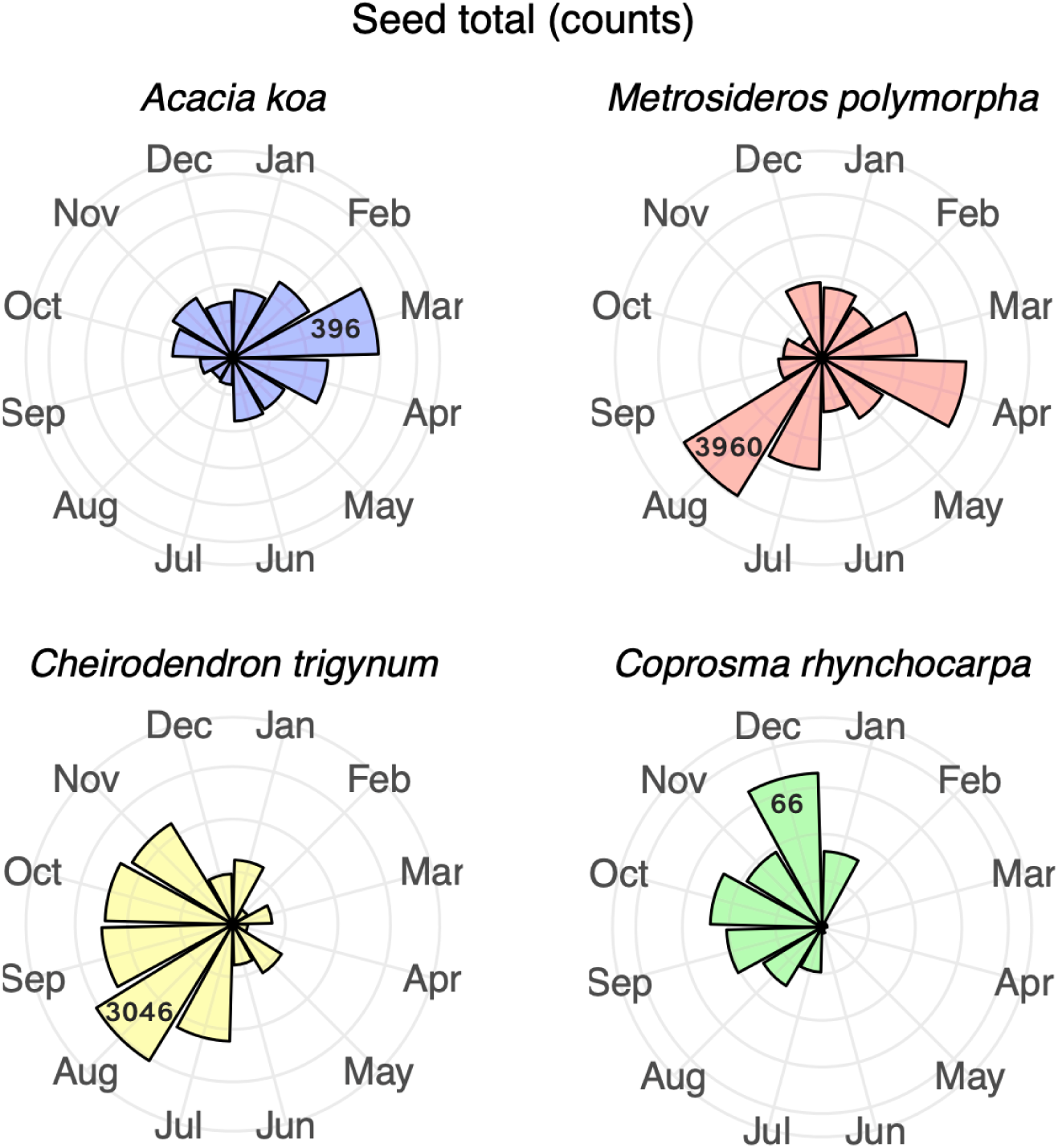
Average monthly seed production of four dominant species in a Hawaiian montane wet forest. Each species is scaled on a different axis length because of large differences in species’ seed production. Maximum seed totals are labeled for each species so relative differences each month can be compared.

All GAM smooth terms (i.e., response curves to climatic factors) were significant (p < 0.001) for seed production by all species (Figure 2 a-d). *A. koa* seed production was the most responsive species to monthly climatic variation, showing increasing seed production with warming temperatures to ∼15 ° C, and increasing seed production with more rainfall and greater PAR up to ∼350 μmol s^−1^m^−2^ (Figure 2a). *C. trigynum* was relatively insensitive to changes in climate compared to the other species with small increases in seed production up to ∼16 ° C, increases up to ∼ 600 mm rainfall, and a flat response to PAR (Figure 2b). Seed production by *C. rhynchocarpa* showed small increases up to ∼16 ° C, decreases in response to rainfall more than ∼500 mm, and rapid decreases in response to PAR above 300 μmol s^−1^m^−2^ (Figure 2c). Seed production from *M. polymorpha* decreased at temperatures above ∼15 ° C, was generally not responsive to seasonal changes in rainfall, and increased slightly with increasing PAR (Figure 2d). Although all climatic smooth terms were significant, the amount of variation in seed production explained by climatic variables was moderate to low depending on species (*A. koa*: R^2^ = 0.22; *C. trigynum*: R^2^ = 0.50; *C. rhynchocarpa*: R^2^ = 0.34; *M. polymorpha*: R^2^ = 0.07).

**Figure 2a-d.**
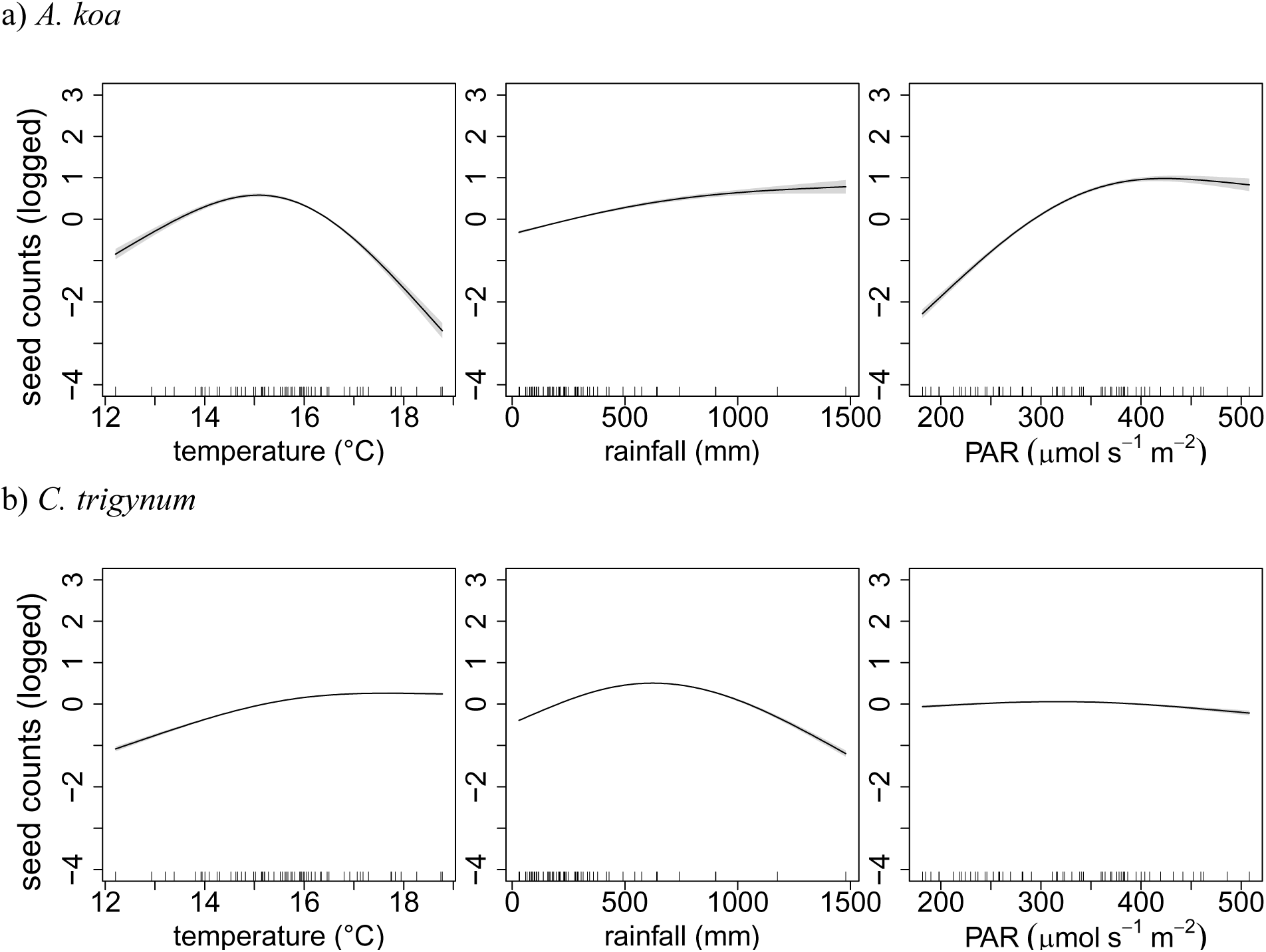
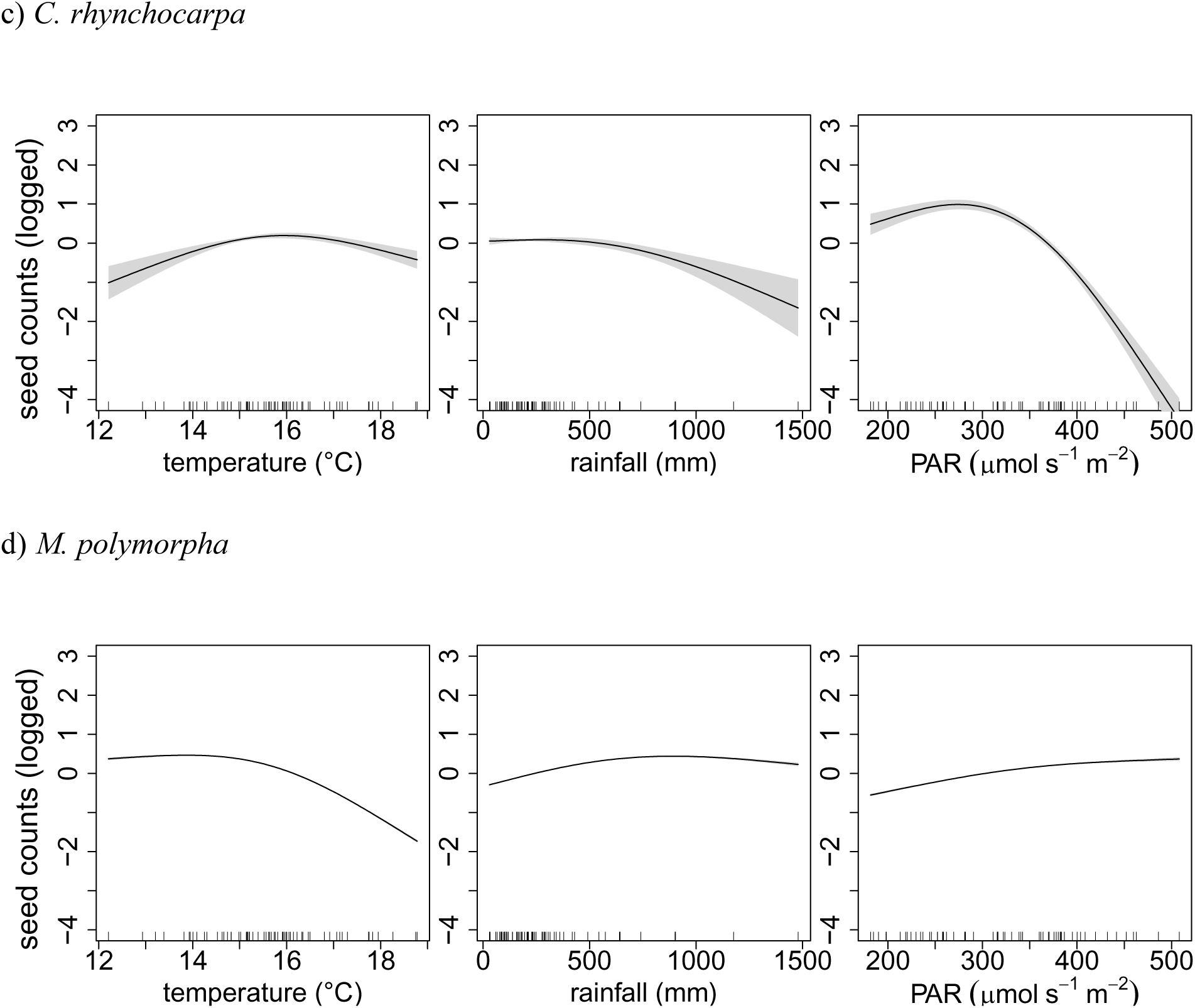
Monthly seed production responses to climatic factors. All GAM smooth terms were significant for all species (p < 0.001; grey shading shows two standard error bounds). (a) *A. koa* (R^2^ = 0.22), (b) *C. trigynum* (R^2^ = 0.50), (c) *C. rhynchocarpa* (R^2^ = 0.34), and (d) *M. polymorpha* (R^2^ = 0.07). Negative log values indicate values between 0-1.

Model comparisons based on AIC showed that the best-fit models for all species included all three climatic factors with close to 100% AIC weight (Table S2a-d).

### leaf litterfall

Leaf litterfall did not show strong seasonality (Figure 3). Only rainfall was significant (p < 0.001) whereas temperature and PAR were not (p > 0.05). Leaf litterfall increased strongly with more rainfall (Figure 4). However the model overall did not explain much variation in leaf litterfall (R^2^ = 0.09). The proportion of variance explained improved when rainfall was the only climatic predictor in the model (R^2^ = 0.14). There were four equivalent best-fit models (ΔAIC <2). The best-fit models each accounting for more than 30% of the AIC weight included either rainfall or PAR separately (Supplementary Table 1e). Two other models considered equivalent best-fit models include rainfall and PAR together, or temperature alone.

**Figure 3.**
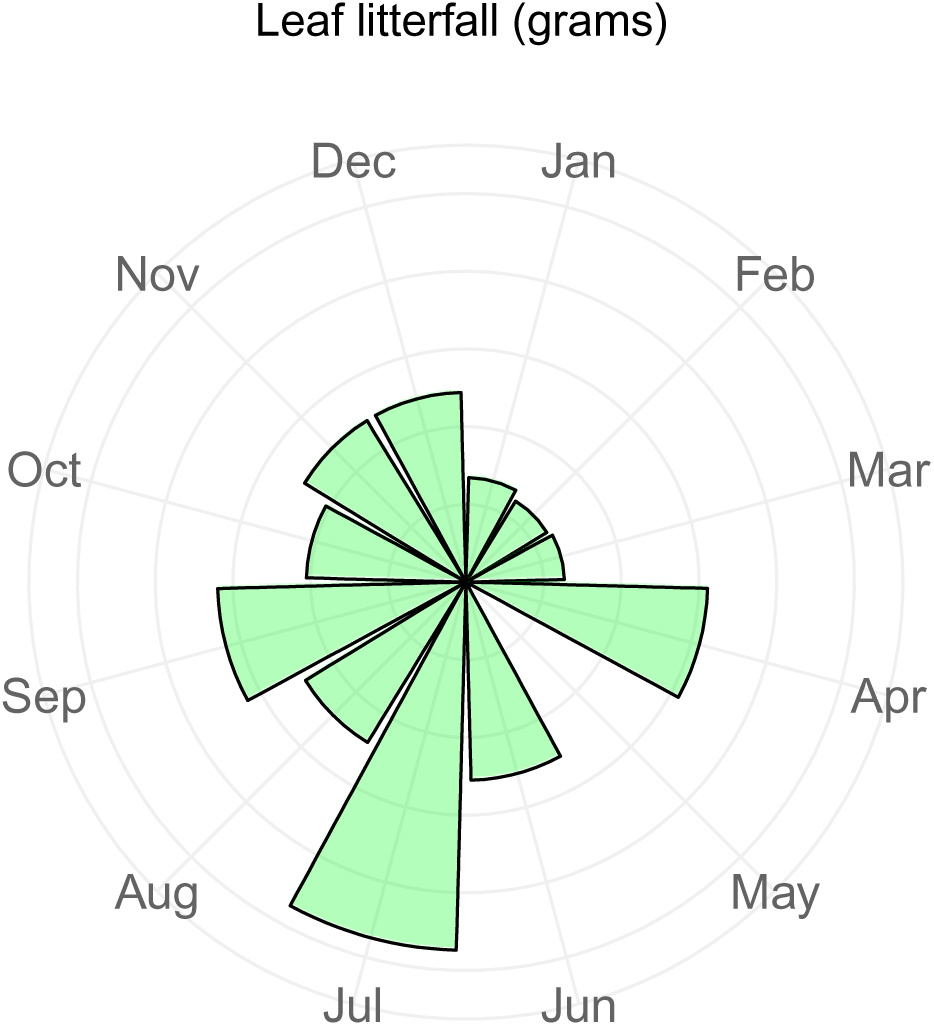
Average monthly community-wide leaf litterfall in a Hawaiian montane wet forest. Grey lines each represent 10 units of leaf litterfall (grams). The month of May averaged 0.7 grams.

**Figure 4.**
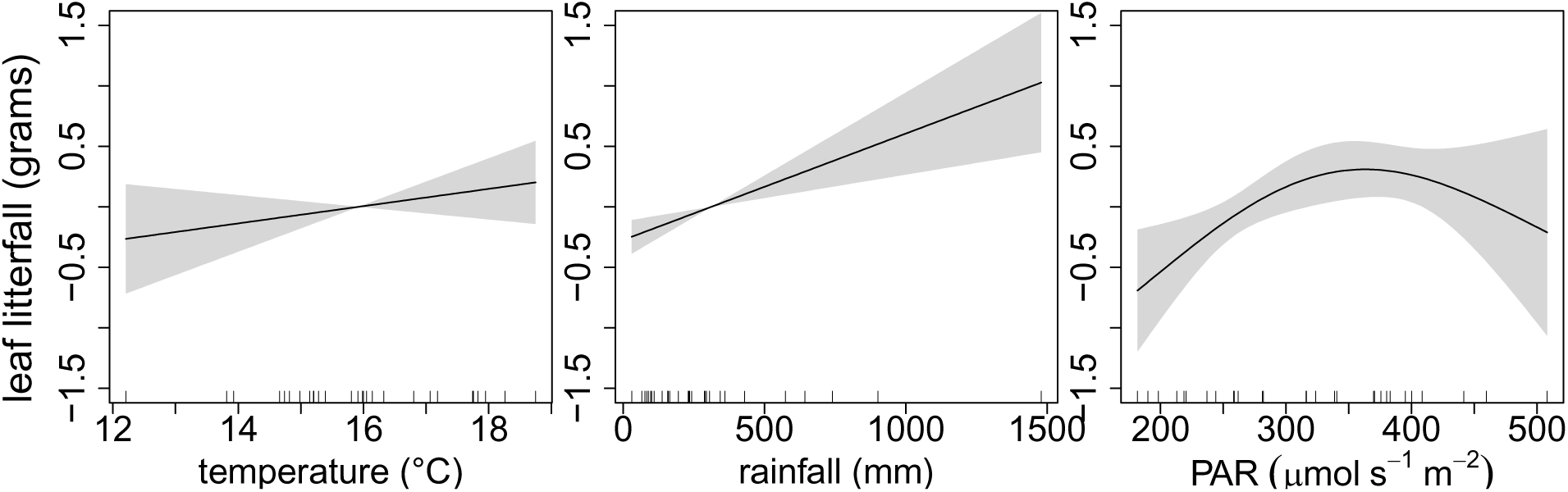
Monthly leaf litterfall responses to climatic factors (R^2^ = 0.09). GAM smooth terms for rainfall was significant (p < 0.001) whereas temperature and PAR were not (p > 0.05; grey shading shows two standard error bounds). Negative log values indicate values between 0-1.

## DISCUSSION

Because the low latitudes are thought to be climatically stable and many species are active year-round, variation in tropical species’ phenologies have been understudied (Cook *et al*. 2012, Chambers *et al*. 2013). However, many tropical plant species are known exhibit distinct phenological patterns (Newstrom *et al*. 1994, Sakai 2001). These patterns may be linked to seasonal changes in the abiotic environment (van Schaik *et al*. 1993). But the degree of sensitivity to climate among tropical species to climatic variations is unknown in many regions, impeding our ability to understand their vulnerability to climate change. We examined seasonal relationships between climate and plant phenology, while accounting for variation due simply to month of the year (as well as changes in daylength; see Supplementary Information). These relationships with climate provide insight into favorable and unfavorable conditions for seed and leaf litterfall production.

### seed production

Response curves showed that seed production was sensitive to changes in climate but no species responded similarly to all three climatic factors. With the exception of one species (*C. trigynum*) seed production was more sensitive to temperature and PAR than to rainfall, supporting our initial hypothesis. Seed production of *A. koa* and *C. rhynocparpa* appeared more sensitive to climatic variation and had higher interannual variability than *M. polymorpha* and *C. trigynum*. Greater interannual variability suggests that photoperiod is not a strong control of reproductive phenology because daylength does not vary each year, yet there was some support for the effect of daylength (Supplementary Information). *M. polymorpha* and *C. trigynum* had weaker responses to climate than *A. koa* and *C. rhynocparpa*. *M. polymorpha*, however, did show a discernable decline with temperatures above 15-16 ° C as did *A. koa*, both foundational species in Hawaiian forests. For all four species at Laupāhoehoe, responses to rainfall and PAR were in the same direction. This could be explained by a large diffuse component of PAR, which can scatter more light through the canopy, as opposed to casting strong shadows, increasing light availability (Roderick *et al*. 2001, Butt *et al*. 2009, 2010, Pau *et al*. 2013). The moderate to low R^2^, particularly for *M. polymorpha*, indicates that factors other than climate play a role in monthly seed production and/or seed production is not very predictable for this species.

An unexamined factor that may explain the timing of seed production and reproductive phenology is conservatism within lineages (Wright & Calderón 1995, Davies *et al*. 2013). A unique feature of Hawaiian plants is that founder populations come from both temperate and tropical regions. Thus phenological patterns may not necessarily indicate local adaptations or be timed to local conditions. Instead they may reflect phylogenetic conservatism from distant ancestors. The four species considered here are all of Australasian descent, with colonists of *A. koa* from Australia and colonists of *M. polymorpha, C. trigynum,* and *C. rhynopc*arpa from New Zealand (Price & Wagner 2018). However the phenologies of their founder populations are difficult to determine in part because these lineages occur in diverse habitats (temperate, arid, tropical dry, tropical wet, etc.) with distinct phenologies.

Although seed production should follow flower production, there may be different climatic cues or resource requirements for seed rather than flower production (Augspurger 1983, Wright & Calderón 2006, Slot & Winter 2016). *A. koa* is a species that is known to flower often but seed development does not always follow. While the presence of flowers was not recorded in censuses, PhenoCam (i.e., a tower-based digital camera; Richardson *et al*., 2009, 2018) observations overlapped with censuses from 2017 and 2018, and seed production followed flowering both years (Fig. S3). Flowering of *A. koa* in PhenoCam images generally occurred December – March (although it began in February in 2018) while seed production peaked, on average, between February – April (Fig. 1).

Even when viable seeds are produced, conditions for establishment may limit recruitment (Inman-Narahari *et al*. 2013). For example, *M. polymorpha* and *C. trigynum* do not appear to be seed or dispersal limited, but instead limited by favorable sites for establishment (Drake 1992, Inman-Narahari *et al*. 2013). In other cases, seed limitation and dispersal failure may contribute to the decline of native species on Hawaii (Chimera & Drake 2010, Inman-Narahari *et al*. 2013).

### leaf litterfall

Community-level leaf litterfall was not well predicted by seasonal changes in climate. Unlike seed production, temperature appeared to have no effect on leaf litterfall. Leaf litterfall was responsive to increasing rainfall. A synthesis of leaf litterfall from tropical South America showed that across sites, litterfall seasonality was associated with rainfall seasonality, wherein sites that had more seasonal litterfall also had more seasonal rainfall and vice-versa (Chave *et al*. 2010) However, there was no relationship between leaf litterfall accumulation and total annual rainfall (Chave *et al*. 2010). Satellite and eddy-covariance measurements have shown positive greening or productivity responses to increases in light availability in tropical wet forests (Huete *et al*. 2006, Saleska *et al*. 2007). In contrast, water-limited sites have shown reduced photosynthesis during the dry season, thus different tropical forest types can exhibit asynchronous responses to climatic variability with water more limiting in dry sites and light more limiting in wet forests (Pau *et al*. 2010, Wu *et al*. 2016, Zhang, Xiao, *et al*. 2016). The dry season greening of tropical forests, even in wet sites, has been intensely debated (Samanta *et al*. 2010, Morton *et al*. 2014) and there are rarely ground measurements of tropical leaf phenology to corroborate satellite measures (Asner & Alencar 2010). One of the few studies that integrated satellite observations and ground-based measures of leaf litterfall in Amazonian forests showed that litterfall was associated with the production of new leaves and greater canopy LAI, which drove increases in satellite measures of greenness (Wu *et al*. 2016). On the contrary, leaf litterfall records from a Panamanian forest appear to coincide with reduced standing leaf area (Detto *et al*. 2018). Given the divergent interpretations of the relationship between leaf litterfall and standing leaf area in tropical forests, identifying site- and species-specific relationships using ground-based observations appear necessary.

### phenological responses to future climate change

The Hawaiian flora is well studied yet there are few systematic investigations of plant phenology and their abiotic drivers. This oversight is significant given the well established patterns of climate change impacts on phenology in other regions (Parmesan & Yohe 2003, Root *et al*. 2003). Phenology is a key axis of functional traits and vital to understanding variation in life history strategies as well as niche differentiation. In Hawaii, a challenge for native plant restoration is the management of invasive species. Identifying differences in the timing of seed production and resource acquisition by native and invasive species, which may change with climate change, should provide insight into mechanisms for establishment and assist in the timing of management efforts (Drake *et al*. 1998, Funk 2013). Additionally, it has been pointed out that species-poor island flora may be accompanied by low functional redundancy, i.e., species perform unique roles in their communities (McConkey & Drake, 2015). Thus communities with strong species’ dependencies are more vulnerable to shifts in the timing of seed or leaf production. Species or populations responding differently in magnitude or direction to climate change may result in phenological mismatches and novel ecological communities (Visser & Both 2005, Thackeray *et al*. 2016).

Shifts in phenology may therefore be viewed as having negative impacts on a community (i.e., phenological mismatch) or viewed positively as an ability to adapt to climate change (Visser & Both 2005). Some hypothesize that species that can track climate change by adjusting their phenologies should persist under a changing climate (Cleland *et al*. 2012). Indeed, research has shown that phenological sensitivity is associated with increased population abundance, however, most of this evidence is from temperate regions where phenological sensitivity is defined as the number of days a species’ shifts a phenological event per degree temperature change (Cleland *et al*. 2012). These species can maintain optimal performance (e.g., flowering or fruiting at a different time), whereas species that do not track climate change may face unfavorable conditions (e.g., climate that is too warm for optimal seed production).

Here we do not examine phenological tracking given the length of our record (∼ 6 years). We show that three of the four dominant species at our site are sensitive to climatic variability, and the direction and magnitude of responses represent favorable or unfavorable periods of growth. End of the century climate projections show an average temperature increase of 2 - 4 °C over the Hawaiian Islands with more warming at high elevations. Increased rainfall and more cloud cover is projected for the windward side of the islands whereas the leeward side is projected to get drier and with fewer clouds (Lauer *et al*. 2013, Zhang *et al*. 2016). As the climate changes in the future, species’ phenologies may shift earlier or later if particular months become unfavorable. Alternatively if species do not shift to more favorable months, fecundity and growth may decline. Ultimately the sensitivity, exposure, and adaptive capacity of species should be considered in determining species’ vulnerabilities to climate change. At least one species (*C. trigynum*), and likely the eight others that reproduced too irregularly for statistical analysis, appear not highly sensitive to climate variability at this site.

Understanding climate change impacts on species and communities has focused on predicting species’ range shifts using bioclimatic envelopes (i.e., correlative models of species presence and mean climate; Guisan & Thuiller 2005, Elith & Leathwick 2009). Shifts in some species’ range and distribution but not others may result in the formation of novel communities (Williams *et al*. 2007). However species’ range limits depend on their ability to complete their life cycle and each species’ distinct reproductive niche is critical to consider (Chuine 2010, Bykova *et al*. 2012). Divergent responses to climate change could alter community composition if reproduction or growth declines for some species but not others. How the reproduction and growth of different species will respond to climate change has potential consequences for future shifts in species distributions and the persistence of biodiversity.

## ACKNOWLEDGMENTS

We thank the National Geographic Committee on Exploration (Grant number #WW-235R-17). Support for HIPPNET has included NSF EPSCoR (Grants No. 0554657 and 0903833) and the USDA Forest Service, the University of Hawai’i, and the University of California, Los Angeles (NSF Grant No. 0546784). We thank Kaikea Blakemore, Christa Nicholas, Leila Ayad, and Evan Blanchard for their help with data collection and processing.

## DATA AVAILABILITY STATEMENT

The data used in this study are archived at the Knowledge Network for Biocomplexity (doi:10.5063/F1QR4VF7).

