## Supporting Information for "Climatic sensitivity of species’ vegetative and reproductive phenology in a Hawaiian montane wet forest"

SEED PRODUCTION AND DAYLENGTH—When daylength was included in models of seed production, all GAM smooth terms (i.e., response curves to temperature, rainfall, PAR, and daylength) were significant ($p < 0.001$) for all species (Fig. S1 a-d). However there was multicollinearity between daylength and temperature (> 0.80 concurvity values, a generalization of collinearity for smooth terms in GAMs). For all species except *C. rhynocarpa*, responsive curves for temperature and other predictors appeared stable with the inclusion of daylength and standard errors were not inflated, indicating that daylength did not interfere with the other predictors (Fig. 2 a-d Main Text). For *C. rhynchocarpa*, the effect of temperature and daylength cannot be determined given the data available. As more data are collected, year-to-year variation in monthly temperatures should allow the independent effects of daylength and temperature to be resolved.

Although all smooth terms were significant, the amount of variation in seed production explained by climatic variables and daylength varied by species (*A. koa*: adj $R^2 = 0.28$; *C. trigynum*: adj $R^2 = 0.47$; *C. rhynchocarpa*: adj $R^2 = 0.56$; *M. polymorpha*: adj $R^2 = 0.05$). Compared to models without daylength (Fig. 2 Main Text), the amount of variation explained slightly increased for *A. koa* and *C. rhynchocarpa*, but slightly decreased for *C. trigynum* and *M. polymorpha*, indicating the overlapping contribution of daylength with other factors, likely temperature given high concurvity values.

Model comparisons of all possible combinations of predictors based on AIC showed that the best-fit models for all species included all three climatic factors and daylength, with close to 100% of the AIC weight (Table S1a-d), for all species except *C. rhynchocarpa*. The best-fit model for *C. rhynchocarpa* included all predictors except for temperature, which had more than 90% of the AIC weight, whereas models without daylength had <1% of AIC weight.

LEAF LITTERFALL AND DAYLENGTH – Only smooth terms for rainfall, daylength, and PAR were significant for community-wide leaf litterfall ($p < 0.001$ for rainfall and daylength; $p < 0.01$ for PAR; $R^2 = 0.30$; Fig. S2). Leaf litterfall increased with more rainfall up to ~ 750 mm, decreased with increasing PAR and increasing daylength, however the responses to rainfall and PAR were not stable when daylength was included. The best-fit model accounting for 96% of the AIC weight included only rainfall (Table S1e and Fig. S2).

Table S1a-e. Model comparisons based on AICc of the full model (all climatic predictors and daylength) and all possible reduced models. Equivalent best-fit models have ΔAIC < 2. All models included 'month' and 'year' smooth terms, as well as an AR(1) error term that accounts for the correlation among observations each month. PAR = photosynthetically active radiation, k = number of parameters, '+' = included in model, 'NA' = excluded in model.

a) *Acacia koa* seed production

| month | year | daylength | PAR | rainfall | temperature | k | AICc | ΔAICc | weight |
|---|---|---|---|---|---|---|---|---|---|
| + | + | + | + | + | + | 13 | 10476.429 | 0.000 | 1.000 |
| + | + | + | + | NA | + | 11 | 10725.075 | 248.646 | 0.000 |
| + | + | NA | + | + | + | 11 | 10756.929 | 280.500 | 0.000 |
| + | + | NA | + | NA | + | 9 | 10845.719 | 369.290 | 0.000 |
| + | + | + | + | + | NA | 11 | 12361.772 | 1885.343 | 0.000 |
| + | + | NA | + | + | NA | 9 | 12449.314 | 1972.886 | 0.000 |
| + | + | + | NA | + | + | 11 | 12787.863 | 2311.434 | 0.000 |
| + | + | + | NA | NA | + | 9 | 12939.737 | 2463.308 | 0.000 |
| + | + | + | + | NA | NA | 9 | 13607.827 | 3131.398 | 0.000 |
| + | + | NA | + | NA | NA | 7 | 13815.968 | 3339.539 | 0.000 |
| + | + | NA | NA | + | + | 9 | 14215.931 | 3739.502 | 0.000 |
| + | + | NA | NA | NA | + | 7 | 15520.320 | 5043.891 | 0.000 |
| + | + | + | NA | + | NA | 9 | 15796.354 | 5319.925 | 0.000 |
| + | + | NA | NA | + | NA | 7 | 16731.953 | 6255.524 | 0.000 |
| + | + | + | NA | NA | NA | 7 | 17100.744 | 6624.316 | 0.000 |
| + | + | NA | NA | NA | NA | 5 | 18435.862 | 7959.433 | 0.000 |

b) *Cheirodendron trigynum* seed production

| month | year | daylength | PAR | rainfall | temperature | k | AICc | ΔAICc | weight |
|---|---|---|---|---|---|---|---|---|---|
| + | + | + | + | + | + | 13 | 36536.361 | 0.000 | 1.000 |
| + | + | + | NA | + | + | 11 | 37067.227 | 530.866 | 0.000 |
| + | + | NA | + | + | + | 11 | 37150.535 | 614.174 | 0.000 |
| + | + | NA | NA | + | + | 9 | 37411.979 | 875.618 | 0.000 |
| + | + | + | + | + | NA | 11 | 38417.596 | 1881.235 | 0.000 |
| + | + | + | NA | + | NA | 9 | 39289.789 | 2753.428 | 0.000 |
| + | + | NA | + | + | NA | 9 | 39670.101 | 3133.740 | 0.000 |
| + | + | NA | NA | + | NA | 7 | 39932.378 | 3396.017 | 0.000 |
| + | + | + | + | NA | + | 11 | 45619.797 | 9083.436 | 0.000 |
| + | + | NA | + | NA | + | 9 | 46298.469 | 9762.108 | 0.000 |
| + | + | + | NA | NA | + | 9 | 46537.053 | 10000.693 | 0.000 |
| + | + | NA | NA | NA | + | 7 | 47518.763 | 10982.402 | 0.000 |
| + | + | + | + | NA | NA | 9 | 48416.931 | 11880.570 | 0.000 |
| + | + | NA | + | NA | NA | 7 | 49535.161 | 12998.800 | 0.000 |
| + | + | + | NA | NA | NA | 7 | 50315.000 | 13778.639 | 0.000 |
| + | + | NA | NA | NA | NA | 5 | 51528.523 | 14992.162 | 0.000 |

c) *Coprosma rhynchocarpa* seed production

| month | year | daylength | PAR | rainall | temperature | k | AICc | ∆AICc | weight |
|---|---|---|---|---|---|---|---|---|---|
| + | + | + | + | + | NA | 11 | 1435.020 | 0.000 | 0.911 |
| + | + | + | + | + | + | 13 | 1439.673 | 4.654 | 0.089 |
| + | + | + | + | NA | NA | 9 | 1479.333 | 44.313 | 0.000 |
| + | + | + | + | NA | + | 11 | 1485.512 | 50.492 | 0.000 |
| + | + | NA | + | NA | + | 9 | 1796.183 | 361.163 | 0.000 |
| + | + | NA | + | + | + | 11 | 1854.781 | 419.761 | 0.000 |
| + | + | NA | + | NA | NA | 7 | 1880.037 | 445.017 | 0.000 |
| + | + | NA | + | + | NA | 9 | 1922.401 | 487.382 | 0.000 |
| + | + | + | NA | + | + | 11 | 2189.128 | 754.109 | 0.000 |
| + | + | + | NA | + | NA | 9 | 2311.017 | 875.997 | 0.000 |
| + | + | + | NA | NA | + | 9 | 2318.998 | 883.978 | 0.000 |
| + | + | + | NA | NA | NA | 7 | 2512.444 | 1077.424 | 0.000 |
| + | + | NA | NA | + | NA | 7 | 2978.284 | 1543.264 | 0.000 |
| + | + | NA | NA | + | + | 9 | 3060.087 | 1625.068 | 0.000 |
| + | + | NA | NA | NA | + | 7 | 3152.733 | 1717.713 | 0.000 |
| + | + | NA | NA | NA | NA | 5 | 3302.961 | 1867.941 | 0.000 |

d) *Metrosideros polymorpha* seed production

| month | year | daylength | PAR | rainfall | temperature | k | AICc | ΔAICc | weight |
|---|---|---|---|---|---|---|---|---|---|
| + | + | + | + | + | + | 13 | 66883.954 | 0.000 | 1.000 |
| + | + | + | NA | + | + | 11 | 67719.508 | 835.554 | 0.000 |
| + | + | NA | + | + | + | 11 | 69402.783 | 2518.829 | 0.000 |
| + | + | + | + | NA | + | 11 | 69856.281 | 2972.327 | 0.000 |
| + | + | NA | NA | + | + | 9 | 70244.011 | 3360.057 | 0.000 |
| + | + | + | NA | NA | + | 9 | 70286.836 | 3402.883 | 0.000 |
| + | + | NA | + | NA | + | 9 | 73655.574 | 6771.620 | 0.000 |
| + | + | NA | NA | NA | + | 7 | 74368.103 | 7484.149 | 0.000 |
| + | + | + | NA | + | NA | 9 | 94198.038 | 27314.084 | 0.000 |
| + | + | + | + | + | NA | 11 | 94951.516 | 28067.562 | 0.000 |
| + | + | + | + | NA | NA | 9 | 95087.803 | 28203.849 | 0.000 |
| + | + | NA | NA | + | NA | 7 | 96310.178 | 29426.224 | 0.000 |
| + | + | + | NA | NA | NA | 7 | 96634.574 | 29750.620 | 0.000 |
| + | + | NA | + | + | NA | 9 | 97581.967 | 30698.013 | 0.000 |
| + | + | NA | + | NA | NA | 7 | 98260.083 | 31376.129 | 0.000 |
| + | + | NA | NA | NA | NA | 5 | 99869.949 | 32985.995 | 0.000 |

e) Community-wide leaf litterfall

| month | year | daylength | PAR | rainfall | temperature | k | AICc | ΔAICc | weight |
|---|---|---|---|---|---|---|---|---|---|
| + | + | NA | NA | + | NA | 8 | 68.819 | 0.000 | 0.965 |
| + | + | + | NA | + | NA | 10 | 76.417 | 7.598 | 0.022 |
| + | + | NA | + | NA | NA | 8 | 77.578 | 8.759 | 0.012 |
| + | + | NA | NA | NA | NA | 6 | 82.064 | 13.245 | 0.001 |
| + | + | + | + | + | NA | 12 | 85.684 | 16.865 | 0.000 |
| + | + | + | NA | + | + | 12 | 86.421 | 17.602 | 0.000 |
| + | + | NA | + | + | + | 12 | 87.534 | 18.715 | 0.000 |
| + | + | + | NA | NA | + | 10 | 89.418 | 20.599 | 0.000 |
| + | + | + | + | NA | + | 12 | 91.269 | 22.450 | 0.000 |
| + | + | + | + | + | + | 14 | 98.355 | 29.536 | 0.000 |

Table S2a-e. Model comparisons based on AICc of the full model and all possible reduced models. Equivalent best-fit models have ΔAIC < 2. All models included 'month' and 'year' smooth terms, as well as an AR(1) error term that accounts for the correlation among observations each month. PAR = photosynthetically active radiation, k = number of parameters, '+' = included in model, 'NA' = excluded in model.

a) *Acacia koa* seed production

| month | year | PAR | rainfall | temperature | k | AICc | ΔAICc | AIC weight |
|---|---|---|---|---|---|---|---|---|
| + | + | + | + | + | 11 | 10756.929 | 0.000 | 1.000 |
| + | + | + | NA | + | 9 | 10845.719 | 88.790 | 0.000 |
| + | + | + | + | NA | 9 | 12449.314 | 1692.385 | 0.000 |
| + | + | + | NA | NA | 7 | 13815.968 | 3059.039 | 0.000 |
| + | + | NA | + | + | 9 | 14215.931 | 3459.002 | 0.000 |
| + | + | NA | NA | + | 7 | 15520.320 | 4763.391 | 0.000 |
| + | + | NA | + | NA | 7 | 16731.953 | 5975.024 | 0.000 |

b) *Cheirodendron trigynum* seed production.

| month | year | PAR | rainfall | temperature | k | AICc | ΔAICc | AIC weight |
|---|---|---|---|---|---|---|---|---|
| + | + | + | + | + | 11 | 37150.535 | 0.000 | 1.000 |
| + | + | NA | + | + | 9 | 37411.979 | 261.444 | 0.000 |
| + | + | + | + | NA | 9 | 39670.101 | 2519.566 | 0.000 |
| + | + | NA | + | NA | 7 | 39932.378 | 2781.843 | 0.000 |
| + | + | + | NA | + | 9 | 46298.469 | 9147.934 | 0.000 |
| + | + | NA | NA | + | 7 | 47518.763 | 10368.228 | 0.000 |
| + | + | + | NA | NA | 7 | 49535.161 | 12384.626 | 0.000 |

c) *Coprosma rhynchocarpa* seed production.

| month | year | PAR | rainfall | temperature | k | AICc | ΔAICc | AIC weight |
|---|---|---|---|---|---|---|---|---|
| + | + | + | + | + | 11 | 37150.535 | 0 | 1.000 |
| + | + | NA | + | + | 9 | 37411.979 | 261.444 | 0.000 |
| + | + | + | + | NA | 9 | 39670.101 | 2519.566 | 0.000 |
| + | + | NA | + | NA | 7 | 39932.378 | 2781.843 | 0.000 |
| + | + | + | NA | + | 9 | 46298.469 | 9147.934 | 0.000 |
| + | + | NA | NA | + | 7 | 47518.763 | 10368.228 | 0.000 |
| + | + | + | NA | NA | 7 | 49535.161 | 12384.626 | 0.000 |

d) *Metrosideros polymorpha* seed production.

| month | year | PAR | rainfall | temperature | k | AICc | ΔAICc | AIC weight |
|---|---|---|---|---|---|---|---|---|
| + | + | + | + | + | 11 | 69402.783 | 0.000 | 1.000 |
| + | + | NA | + | + | 9 | 70244.011 | 841.228 | 0.000 |
| + | + | + | NA | + | 9 | 73655.574 | 4252.791 | 0.000 |
| + | + | NA | NA | + | 7 | 74368.103 | 4965.321 | 0.000 |
| + | + | NA | + | NA | 7 | 96310.178 | 26907.395 | 0.000 |
| + | + | + | + | NA | 9 | 97581.967 | 28179.184 | 0.000 |
| + | + | + | NA | NA | 7 | 98260.083 | 28857.301 | 0.000 |
| + | + | NA | NA | NA | 5 | 99869.949 | 30467.166 | 0.000 |

e) Community-wide leaf litterfall.

| month | year | PAR | rainfall | temperature | k | AICc | ΔAICc | AIC weight |
|---|---|---|---|---|---|---|---|---|
| + | + | NA | + | NA | 6 | 73.170 | 0.000 | 0.315 |
| + | + | + | NA | NA | 6 | 73.214 | 0.045 | 0.309 |
| + | + | NA | NA | + | 6 | 74.347 | 1.177 | 0.175 |
| + | + | + | + | NA | 8 | 74.648 | 1.478 | 0.151 |
| + | + | NA | + | + | 8 | 78.242 | 5.073 | 0.025 |
| + | + | NA | NA | NA | 4 | 79.517 | 6.347 | 0.013 |
| + | + | + | NA | + | 8 | 80.160 | 6.990 | 0.010 |
| + | + | + | + | + | 10 | 82.897 | 9.727 | 0.002 |

Figure S1. Monthly seed production responses to daylength and climatic factors. All GAM smooth terms were significant for all species ($p < 0.001$; grey shading shows two standard error bounds). (a) *A. koa* ($R^2 = 0.22$), (b) *C. trigynum* ($R^2 = 0.50$), (c) *C. rhynchocarpa* ($R^2 = 0.34$), and (d) *M. polymorpha* ($R^2 = 0.07$). Negative log values indicate values between 0-1.

a) *A. koa*

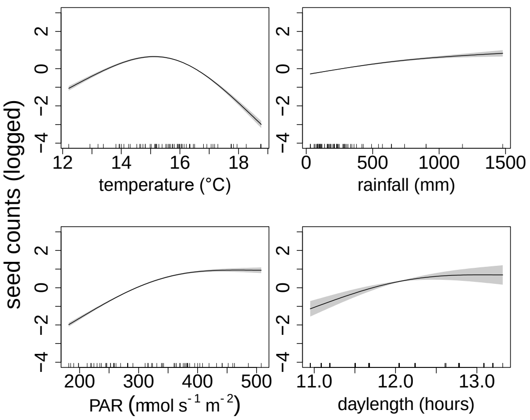


b) *C. trigynum*

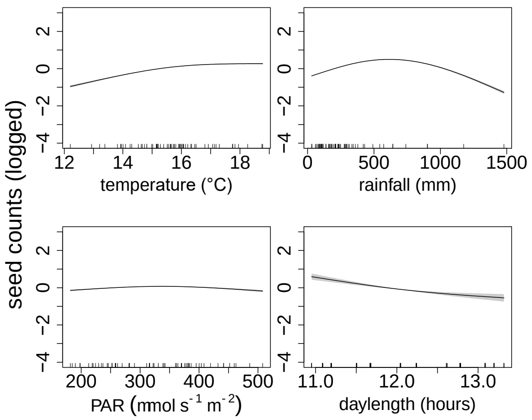


c) *C. rhynchocarpa*

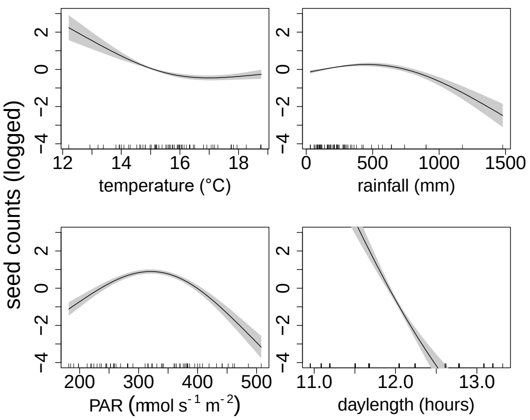


d) *M. polymorpha*

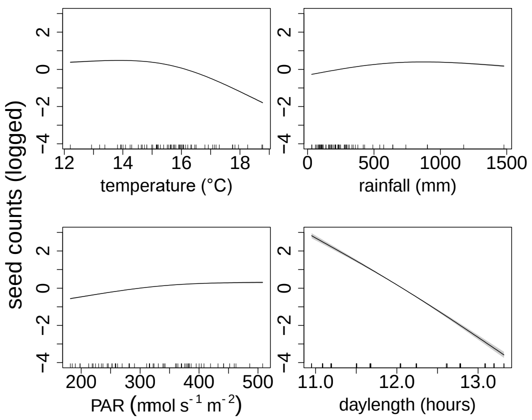

Figure S2. Monthly leaf litterfall responses to daylength and climatic factors. Smooth terms for rainfall, PAR, and daylength were significant for community-wide leaf litterfall ($p < 0.001$ for rainfall and daylength; $p < 0.01$ for PAR) whereas temperature was not significant ($p > 0.05$; $R^2 = 0.30$; grey shading shows two standard error bounds). Negative log values indicate values between 0-1.

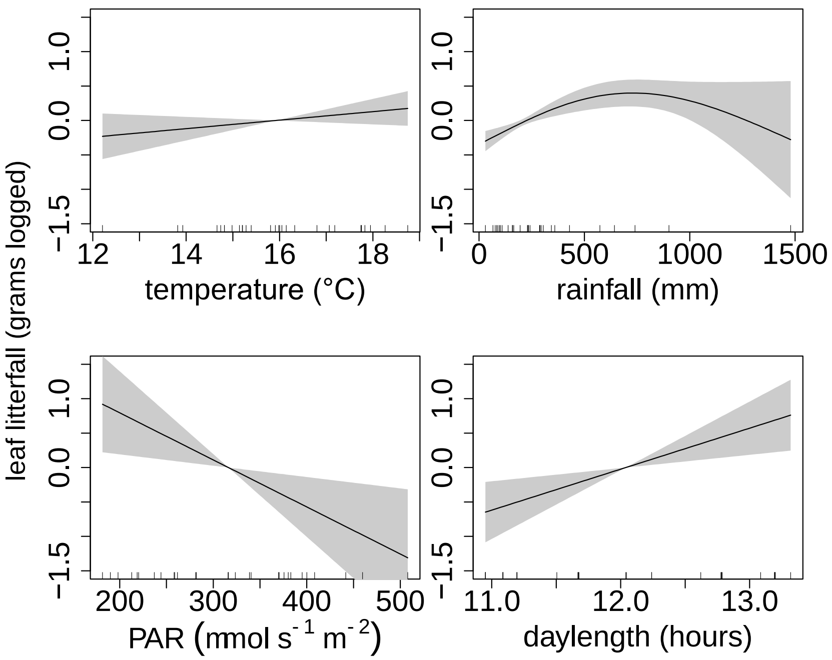

Figure S3. Daily images of four *Acacia koa* canopies from November 2016 to March 2018 were examined for the presence (yes or no) of inflorescences and pods (although no litter basket censuses were collected for 2016 and part of 2017).

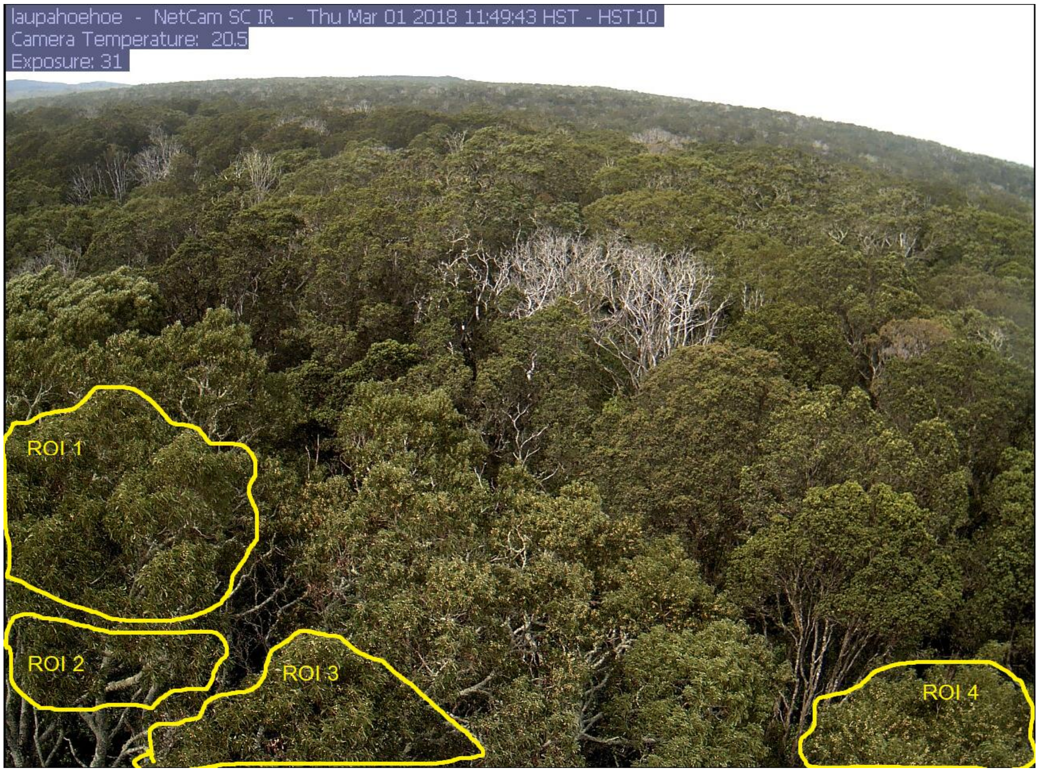
